# Social integration predicts mitochondrial DNA copy number in rhesus macaques

**DOI:** 10.1101/408849

**Authors:** Reena Debray, Noah Snyder-Mackler, Jordan Kohn, Mark Wilson, Luis Barreiro, Jenny Tung

## Abstract

In many social mammals, social adversity predicts compromised health and reduced fitness. These effects are thought to be driven in part by chronic social stress, but their molecular underpinnings are not well understood. Recent work suggests that chronic stress can affect mitochondrial copy number, heteroplasmy rates, and function. Here, we tested the first two possibilities, for the first time in nonhuman primates. We manipulated dominance rank in captive female rhesus macaques (n=45), where low rank induces chronic social stress, and measured mitochondrial DNA copy number and heteroplasmy in five peripheral blood mononuclear cell types from each study subject. We found no effect of dominance rank on either mtDNA copy number or heteroplasmy rates. However, grooming rates, a measure of affiliative social behavior predicted by high social status, was positively associated with mtDNA copy number in B cells, cytotoxic T cells, and monocytes. Our results suggest that social interactions can influence mtDNA regulation in immune cells. Further, they indicate the importance of considering both affiliative and competitive interactions in investigating this relationship.

## Introduction

In social mammals, the rate and outcome of competitive and affiliative interactions often co-vary with fitness-related traits (1,2). These relationships arise in part because social interactions influence access to other fitness-determining factors, such as food or mates, and in part because individual condition can shape social interactions. However, recent evidence also points to a third explanation: direct effects of social interactions on animal physiology, often in connection to socially induced stress (3). Socially subordinate or isolated individuals exhibit elevated glucocorticoid levels (4), up-regulated beta-adrenergic signaling (5), and altered gene regulation (6), which in turn may contribute to social gradients in disease and mortality rates (7,8).

Recent evidence indicates that social adversity may also impact mitochondrial DNA (mtDNA) content, function, and regulation. Experimental studies in rodents indicate that chronic social stress reduces mitochondrial energetic capacity, alters mtDNA gene expression, and changes the complement of mitochondria-associated proteins and metabolites (9). Further, cortisol treatment induces glucocorticoid receptor binding to mtDNA (10), suggesting a link between hypothalamic-pituitary-adrenal axis regulation of stress and mitochondrial activity. Depression in women and physical stressors in mice also predict higher mtDNA copy number and heteroplasmy (mtDNA genetic variation within an individual) in blood and saliva (11,12). Together, these findings suggest that mtDNA biology is sensitive to chronic stress, including social stress. However, this hypothesis has not been tested in nonhuman primates, despite extensive work on the consequences of social interactions for other measures of physiology, health, and fitness (1,2,13).

Here, we address this gap by studying captive female rhesus macaques (*Macaca mulatta*), in which social status (i.e. dominance rank) was experimentally manipulated via controlled introduction of females into new social groups (14). Female rhesus macaques form stable, linear dominance hierarchies, and social status affects access to resources, social control, and exposure to psychosocial stress (15). In our experimental population, later introduction predicted lower rank, which in turn predicted higher rates of received harassment and lower rates of affiliative grooming behavior (14). To test the effects of social interactions on mtDNA biology, we measured mtDNA copy number and heteroplasmy in five purified peripheral blood mononuclear cell (PBMC) types. Motivated by previous observations in humans and mice (11,12), we predicted that social subordination and relative social isolation—that is, increased exposure to social stressors—would lead to higher mtDNA copy number and heteroplasmy rates.

## Materials and Methods

### a. Behavioral data collection

We studied nine social groups, each composed of five unrelated adult female rhesus macaques (described in prior work (14): Table S1). Females were sequentially introduced such that later introduction predicted lower dominance rank. Rank was quantified based on agonistic behavioral interactions obtained during focal sampling (16) (121.5 hours of observation), using Elo ratings (17). To measure affiliative behavior, we used the proportion of focal observation time that a female spent grooming or being groomed.

### b. mtDNA copy number quantification

DNA used in this study was obtained from previous work (14). In brief, PBMCs were purified from 12-20 mL of blood from each female, five PBMC cell types (B cells, cytotoxic T cells, helper T cells, monocytes, and natural killer cells) were sorted into separate populations, and RNA and DNA were extracted.

We measured mtDNA copy number in duplicate using quantitative PCR (qPCR), targeting a region of the mitochondrial genome and a single-copy region of the nuclear genome (n=213 samples for which sufficient DNA was available: see also Text S1). Using the cycle threshold (Ct) values of the qPCR runs, we calculated mtDNA copy number as 2^(mean Ct of nuclear DNA replicates – mean Ct of mtDNA replicates)^, and log-transformed mtDNA copy number for downstream analysis. Our mtDNA copy number estimates were robust to use of alternative nuclear and mtDNA loci (Figure S1).

We modeled natural log-transformed mtDNA copy number as a function of social interactions (dominance rank or grooming behavior), controlling for age, cell type, and qPCR plate/batch (Text S2), using linear mixed effects models. We first investigated social interaction effects on mtDNA copy number across all cell types. Next, to test for heterogeneity in the effects of social interactions across cell types, we modeled each cell type separately and used a Bayesian meta-analytic approach to investigate shared effects across cell types (18).

### c. Heteroplasmic variants in mtDNA

We quantified mitochondrial heteroplasmy using mtDNA-mapped reads from previously generated, cell type-specific RNA-seq data for each study subject (14) (accession number GSE83307; see also Text S3) and the program mitoCaller (19). To account for sequencing coverage, we removed mtDNA positions with fewer than 200 reads and randomly subsampled sites with >200 reads down to exactly 200 reads. Following (12), we classified a site as heteroplasmic if the minor allele frequency was at least 4% within a sample.

We modeled heteroplasmy using a binomial mixed model approach (20), controlling for age and cell type. Our response variable was the number of heteroplasmic sites in the mtDNA genome relative to the number of analyzable sites for each sample.

## Results

### a. Grooming behavior, but not dominance rank, predicts mtDNA copy number

Dominance rank did not predict mtDNA copy number across all cell types together (ß=–0.033, *p=*0.931, Table 1) or in any individual cell type (all p>0.1, Table 1). However, mtDNA copy number was positively associated with the amount of time females spent grooming (ß=0.017, *p=*0.038, Table 2). This association varied across cell types, with the strongest effect in B cells and a nonsignificant trend in monocytes and cytotoxic T cells (Figure 1). Consistent with these observations, formal meta-analysis strongly supported a shared positive effect of grooming on mtDNA copy number in B cells, monocytes, and cytotoxic T cells (Bayes factor=15.67). As expected (21), copy number varied systematically across cell types (Figure S2) and tended to decrease with age (Figure S3).

**Table 1.** Effects of dominance rank and age on mtDNA copy number.

|  | Coefficient (B) | Standard error | Test statistic | p-value |
| --- | --- | --- | --- | --- |
| **<u>All cell types</u>** |  |  |  |  |
| Dominance rank | -0.033 | 0.038 | -0.087 | 0.931 |
| Age | -0.017 | 0.011 | -1.660 | 0.099 |
| **<u>Monocytes</u>** |  |  |  |  |
| Dominance rank | -0.010 | 0.073 | -0.138 | 0.891 |
| Age | 0.016 | 0.019 | 0.822 | 0.420 |
| **<u>Natural killer cells</u>** |  |  |  |  |
| Dominance rank | -0.048 | 0.098 | -0.491 | 0.626 |
| Age | 0.014 | 0.027 | 0.512 | 0.611 |
| **<u>B cells</u>** |  |  |  |  |
| Dominance rank | 0.036 | 0.074 | 0.486 | 0.630 |
| Age | -0.032 | 0.020 | -1.633 | 0.110 |
| **<u>Helper T cells</u>** |  |  |  |  |
| Dominance rank | -0.064 | 0.095 | -0.687 | 0.496 |
| Age | **-0.056** | **0.026** | **-2.110** | **0.041** |
| **<u>Cytotoxic T cells</u>** |  |  |  |  |
| Dominance rank | 0.059 | 0.078 | 0.768 | 0.447 |
| Age | **-0.052** | **0.024** | **-2.158** | **0.037** |

**Table 2.**
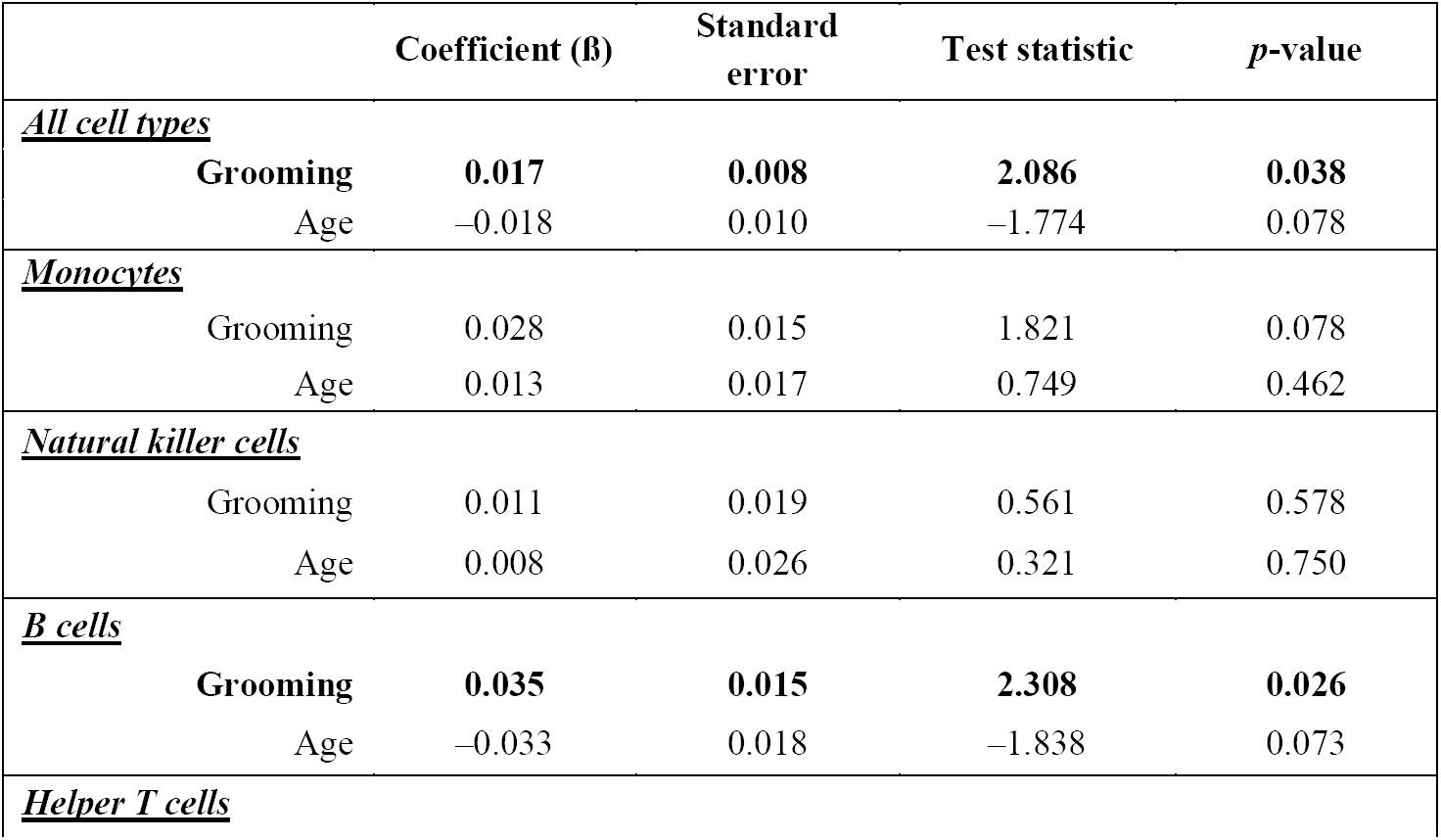
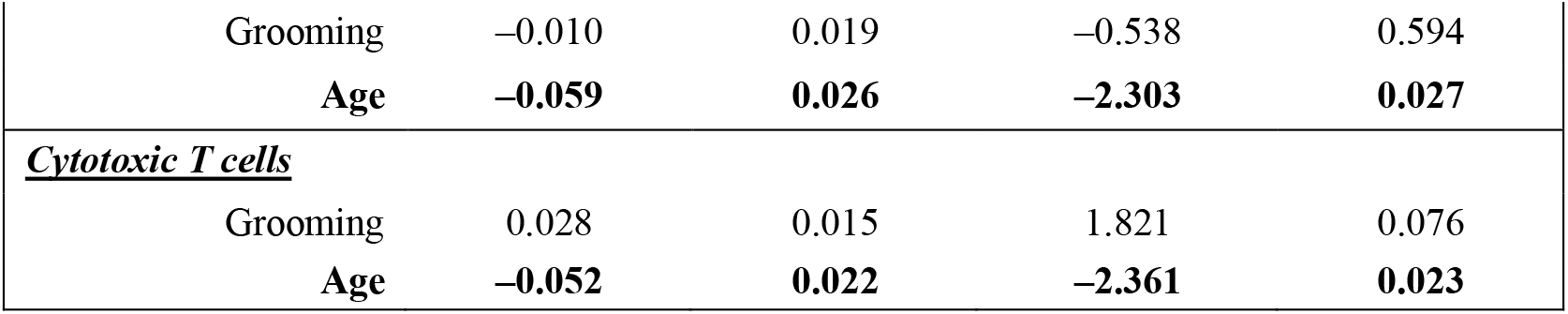
Effects of grooming and age on mtDNA copy number.

**Figure 1.**
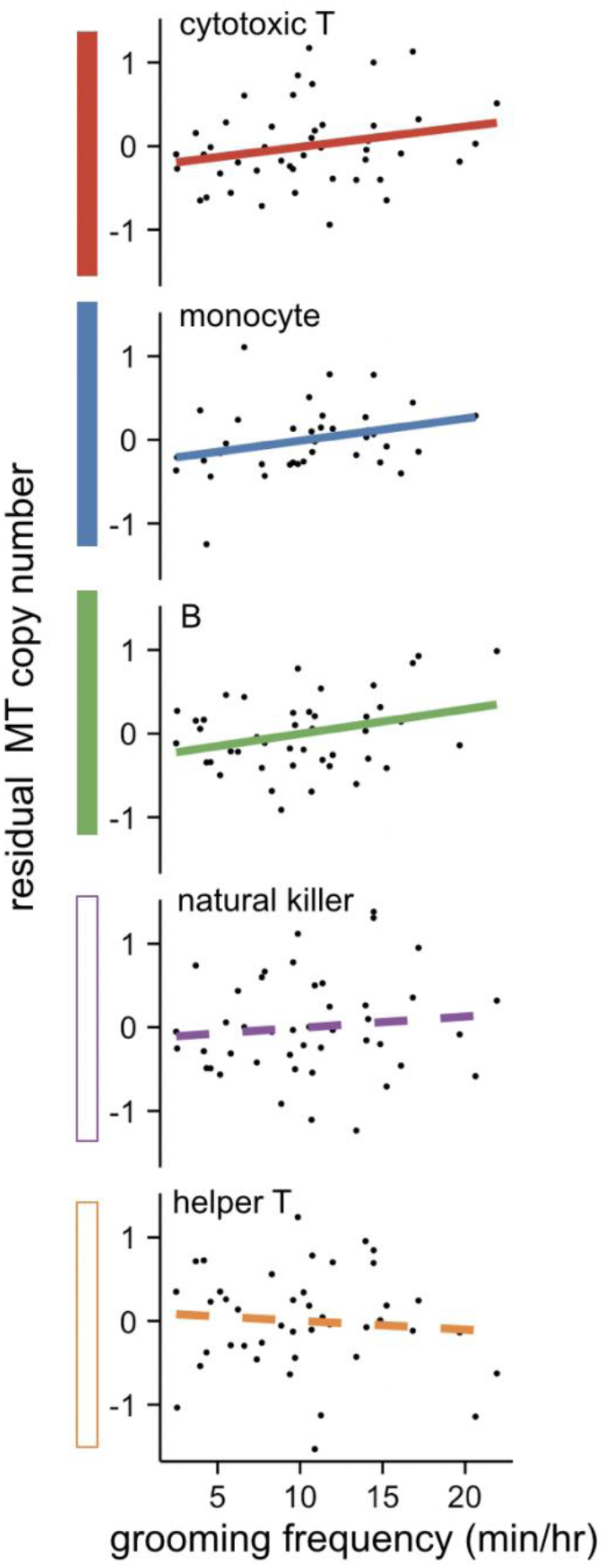
Cell type-specific effects of grooming on mtDNA copy number. Shaded boxes (left) show shared positive effects of grooming on mtDNA copy number based on meta-analysis (empty boxes = no effect). For visualization purposes, the y-axis shows the residuals of a model controlling for age and qPCR batch.

### b. Social interactions do not predict heteroplasmy

Consistent with previous findings in humans (19), heteroplasmic sites occurred at low levels in each sample (mean number of heteroplasmic sites = 5.93 ± 4.10 s.d.). Most heteroplasmic mutations were private within samples, though samples from the same individual tended to share heteroplasmic sites (Figure S4). Neither dominance rank nor grooming rates were significant predictors of overall heteroplasmy rates (all p>0.1, Tables S4, S5, Figure 2).

**Figure 2.**
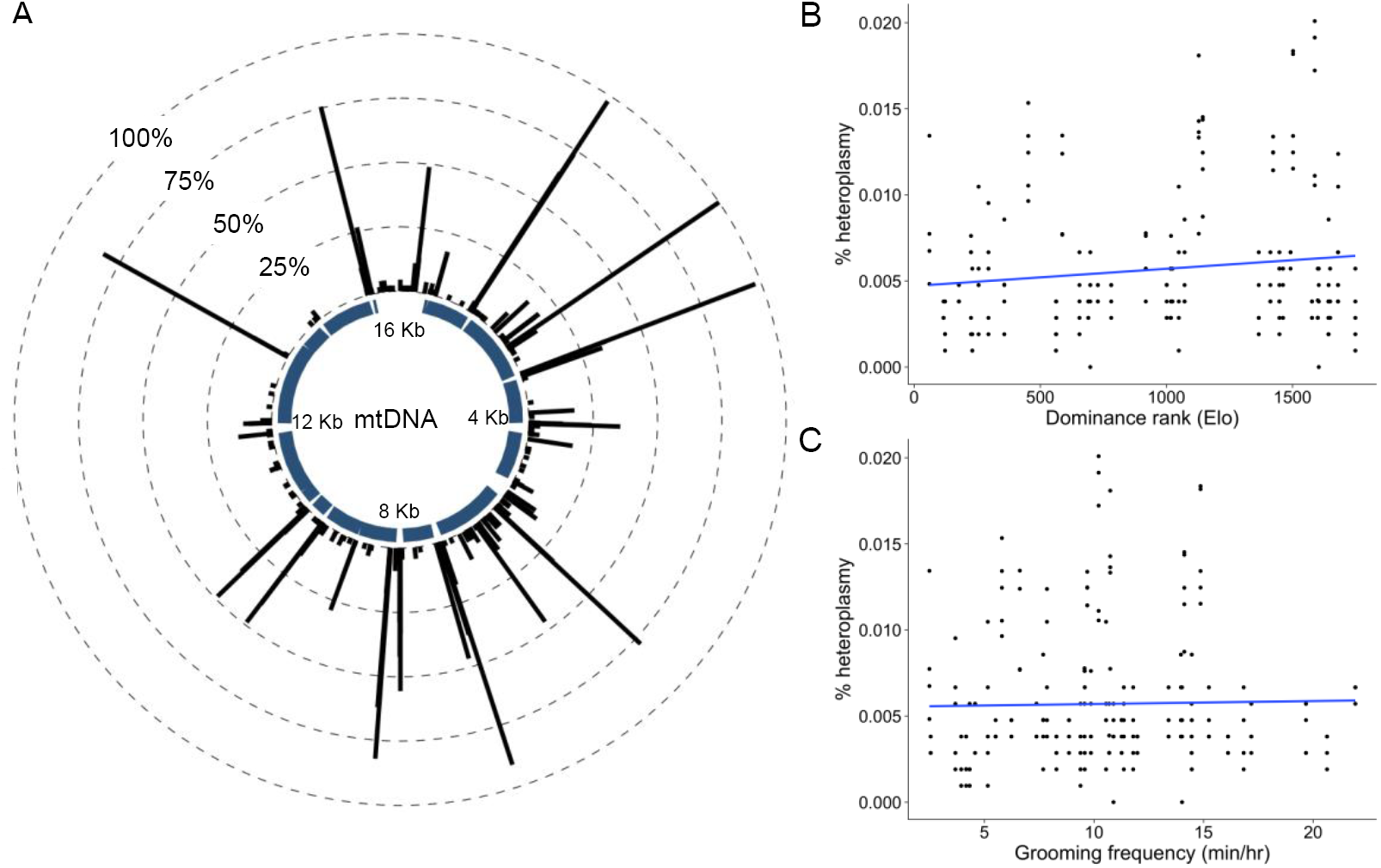
No effect of dominance rank or grooming on mtDNA heteroplasmy. (a) Locations and frequencies of heteroplasmic sites. The *θ*-axis shows mtDNA coordinates based on (22). The y-axis shows the number of individuals (out of 45) that carried a heteroplasmy in at least one cell type. (b,c) Neither dominance rank (b) nor grooming behaviour (c) predicts heteroplasmy rates, controlling for age and cell type.

## Discussion

In contrast to our predictions, we found no effect of social status on mtDNA copy number. Further, high social affiliation was positively, not negatively, predictive of mtDNA copy number. Thus, our results support the idea that social interactions can influence mitochondrial biology, but they do so here in a surprising direction. One potential explanation for this difference is that previous studies focused on heterogeneous cell populations (11,12), whereas here we analyzed purified populations. mtDNA copy number varies among cell types (21) (Figure S2). Thus, social factors that influence cellular composition could also affect apparent differences in copy number. Alternatively, low social status may affect mitochondrial biology differently than previously studied factors, such as early childhood adversity (23) or depression (11). We note, however, that our results are broadly consistent with findings that mtDNA copy number in the rat hippocampus and striatum was reduced by daily exposure to corticosterone and chronic exposure to mild physical stressors (24).

Together, our findings extend experimental studies of social factors on mitochondrial biology to primates, and add important data on females (10). We do not yet understand the importance of grooming effects on mtDNA copy number for cellular and organismal function. Increases in copy number may signify increased energetic potential or compensate for loss of mitochondrial function when some mtDNA copies are mutated. Additionally, while mtDNA copy number has been linked to many disorders, the directionality of this relationship depends on the disorder. For example, mtDNA copy number in blood is positively associated with rates of mitochondrial encephalomyopathies (26) but negatively associated with rates of metabolic syndrome (27).

Because this study used previously extracted DNA samples, we were not able to measure direct indicators of mitochondrial function. Future studies that measure both mtDNA copy number and mitochondrial metabolism in the same samples will clarify how mitochondrial quantity and quality combine to respond to social environmental cues. Such studies have the potential to shed new light on the relationship between mtDNA biology, human health, and its importance for Darwinian fitness in social mammals more broadly.

## Acknowledgments

We thank J. Whitley, A. Tripp, N. Brutto, and J. Johnson for collecting behavioral data, N. Cai for advice on quantifying mtDNA copy number, and T. Reddy for access to the qPCR thermocycler.

## Author contributions

JT, LBB, MEW, NSM, and RRD designed the study. RRD, NSM, and JK collected the data. RRD, NSM, and JT analyzed the data and wrote the paper, with contributions from all authors.

## Data accessibility

Data are included as supplementary material, in SRA (accession number GSE83307), and at https://github.com/reenadebray/mtDNA_copy_number.

